# A GSVA based gene set synergizing with CD4+T cell bearing harmful factors yield risk signals in HBV related diseases via amalgamation of artificial intelligence

**DOI:** 10.1101/2022.01.19.476726

**Authors:** Jun Huang, Chunbei Zhao, Xinhe Zhang, Qiaohui Zhao, Yanting Zhang, Liping Chen, Guifu Dai

**Affiliations:** School of Life Sciences, Zhengzhou University, Zhengzhou City 450001,Henan province,P.R. China; Department of Gastroenterology and Hepatology, Key Laboratory of Gastroenterology and Hepatology, Ministry of Health, State Key Laboratory for Oncogenes and Related Genes, Shanghai Institute of Digestive Disease, Renji Hospital, School of Medicine, Shanghai Jiaotong University, Shanghai 200001, P.R. China

**Author notes:** Correspondence should be addressed to: Dr. Jun Huang,., Dr. Guifu Dai,; School of Life Sciences, Zhengzhou University, No.100 Science Avenue, Zhengzhou City 450001, Henan Province P.R. China.

**Keywords:** HBV related diseases, CLST, infiltrating immune cells, CD4+T cells, risk signal

## Abstract

Genes encoding chemokines and extracellular matrix (ECM) play pivotal roles in chronic HBV infection (CHB), HBV related fibrosis (HBV-LF) and hepatocellular carcinoma (HBV-HCC). The landscape and potential of these genes in prognosis across diseases stages have not been fully and systemically understood. In this study, we defined an HBV-LF associated gene set comprised of chemokines and ECM related genes directly induced by initial HBV infection through GSVA algorithm that named as CLST (C stands for CXCL9, CXCL10, CCL19 and CCL20; L for LUM; S for SOX9 and SPP1; T for THBS1, THBS2) and evaluated its biomarker values in CHB and HBV-LF. Enrichment scores (ES) of CLST was subsequently observed synergized with activated CD4+T cells (aCD4) highly related to T helper cell 17 (TH17) associated genes and immune checkpoints and addressed as risk signals due to bearing harmful prognosis factors in tumor tissues of patients with HBV-HCC. Dual higher enrichment score (ES) of CLST and aCD4 in HBV-HCC patients exhibited worse overall survival (OS). Feature genes specific to these two gene sets showed promising clinical relevance in early-stage of HBV-HCC definition and OS prediction incorporating laboratory parameters via artificial intelligence (AI) systems. Finally, a novel mechanistic insight into the issue was proposed that PEG IFN-α as an immunotherapy through modulating CLST signal in treatment responders and these immune signals down-regulation could be beneficial for HBV related diseases control and prevention. Together, our study provides GSVA and AI derived immunogenomic prognosis signatures and clinical utility of these signals will be benefit for HBV related diseases cure.

## Introduction

Chronic hepatitis B infection (CHB) is one of the leading causes for HBV-LF and highly associated with malignant liver diseases such as HBV-HCC. HBV-LF induced by CHB refers to abnormal proliferation of connective tissue in the liver, characterized by excessive precipitation of diffuse extracellular matrix in normal region(1-3). In severe cases, it develops to cirrhosis and eventually resulted in liver failure or HBV-HCC(4). According to the previous reports, the incidence of HCC in CHB patients is about 3% per 1000 person-years(5), which brings heavy economic pressure and psychological burden to many families. Overall, HBV related diseases have been global public health issues, especially in the Asia-Pacific region where HBV is highly prevalent(6-9).

First-line anti-HBV drugs approved by FDA including PEG IFN-α and nucleoside (acid) analogs (NAs) are not yet effective in achieving functional cure referring to hepatitis B surface antigen (HBsAg) and covalently closed circular DNA (cccDNA) elimination (10-13). Progressive LF were recently reported in CHB patients during long-term nucleoside (acid) analogs treatment (14). The response rate of PEG IFN-α therapy in CHB patients is about 30% and higher than NAs but the underlying antiviral mechanism remains elusive. It is an urgent to uncover factors involved in efficient PEG IFN-α therapy and R&D of cure medicine to solve long-term viremia (LLV) and drug resistance dilemma that NAs may face in the future (15-17). Discovery of innovative therapeutic targets is inseparable from an in-depth understanding of virus-host interaction. Investigating and confirming targets that directly blocking viral replication or indirectly through regressing HBV mediated pathogenesis from more than 20,000 protein coding genes is a huge project, previously. With the development of gene microarray and high-throughput RNA-sequencing, a large amount of high-quality gene expression date were generated (6, 18-23) and majority of these data uploaded to web-accessible databases give rise to the possibilities to carry out integrated bioinformatics analysis (6, 24-29).

The orchestra of viral factors, parenchymal hepatic cells, tissue-resident lymphocytes and extrahepatic immune cells in liver microenvironment is highly associated with HBV related diseases processes (30).Until now, intrahepatic landscape of immunophenotypes and their diagnosis values remind largely unknown and few studies identified core immune signals during disease progression from initial HBV infection to HBV-HCC by Single-sample gene set enrichment analysis (ssGSEA) widely used in immunogenomic biosignatures identification (3, 29, 31-33). Intrahepatic chemokines play pivotal roles in the initiate recruiting extrahepatic pathogenic T cells (34-37) (38), ECM deposition are drivers of malignant liver diseases (39, 40), but these genes in liver tissues as a whole and their cross-talking with intrahepatic immune cells across disease stages for candidate biomarkers development,as well as therapeutical targets for immune interruption are still far from being investigated.

Although several models based on clinical features, laboratory parameters and gene expression by performing AI algorithms were constructed for HCC tumor detection and survival prediction (41-45), few take maker genes belonging to immune gene sets into consideration and the optimal model with promising prognostic value is far from reaching a general consensus(44-46). In this study, we performed immunogenomic profiling aiming to uncover core immune signal involved in all stages of HBV related diseases(initial HBV infection, CHB, HBV-LF and HBV-HCC). The potential roles of core immune signal in recruiting infiltrating lymphocytes, predicting liver injury, detecting liver samples with HBV-LF and as risk survival factor in HBV-HCC were also addressed. Then, powerful AI systems were incorporated to construct prognostic models for early stage of HBV-HCC identification and OS prediction. Finally, the influence of standard IFN-α therapy on core immune signal in responder at CHB phases was evaluated to validate our exploration. In this study, a new strategy and workflow for novel biomarkers identification based on GSVA and AI algorithms through analyzing components of immunogenomic gene set was established that will be helpful for anti-HBV immunotherapies exploration and clinical-decision making.

## Materials and methods

### Raw data collection and proceeding

12 GEO datasets including GSE84044 (HBV-LF),GSE83148 (CHB),GSE65359 (CHB),GSE52752 (HBV infected human hepatocyte chimeric mice, HBV-mice), GSE69590 (HBV infected primary human hepatocytes, HBV-PHH),GSE138569 (HBV-PHH), GSE62232 (HBV-HCC),GSE94660 (HBV-HCC),GSE14520 (HBV-HCC),GSE121248 (HBV-HCC),GSE25097 (cirrhotic, HCC) and GSE66698 (PEG IFN-α treatment responders) were available on Gene Expression Omnibus(GEO) database. CHCC-HBV consisted of patients with HBV-HCC was collected from NODE (https://www.biosino.org/node). R Studio (Version 1.4.1103) were used to raw data proceeding (normalization, gene ID convention, clinical information collection) based on recommend R packages.

### Identification of DEGs

DEGs (S1/S0, S2/S0, S3/S0, S4/S0) of GSE84044 were downloaded from supplementary material of the previous study (6) and visualized via Numbers, an APP of apple.inc. DEGs (G1/G0, G2/G0, G3/G0, G4/G0) of GSE84044 were screened primarily via “Limma” R package and visualized via “ggplot2” R package, “pheatmap” R package or “EnhancedVolcano” R package. As for “EnhancedVolcano” R package, up-regulated genes with fold change (FC) >1.5 and *p* value<0.05 were considered statistically significant. Venn analysis were used to investigate common overlapping DEGs.

### Functional annotation of DEGs and hub genes screening

GO analysis were performed to investigate the biological function annotation of 64 DEGs of GSE84044 using online database (DAVID Bioinformatics Resources 6.8) and visualized via “ggplot2” R package. KEGG signaling pathway analysis were based on online database (DAVID Bioinformatics Resources 6.8) and also visualized via “ggplot2” R package. A PPI network of 64 DEGs from GSE84044 containing 57 nodes and 89 edges was conducted on STRING database. Cytoscape software was utilized to visualize and further screen hub genes. Finally, protein and protein interaction(PPI) analysis of member genes of aCD4 were conducted and visualized using online tools at STRING database.

### The ssGSEA score calculation

The ssGSEA scores of 28 liver infiltrating lymphocytes (LILs) and CLST signal in each sample from microarray data of GEO database or CHCC-HBV were calculated primarily via “GSVA” R package according to the previous studies(31, 32). In detail, 28 gene sets consisted of cell specific marker genes represent 28 LILs (31) and CLST consisted of hub genes defined by us in this study according to previous studies(3, 33).

### Correlation and comparison

“Hmisc” R package was utilized to calculate the correlations between selected gene and LILs. R package “ggcorrplot” was utilized to calculate the correlations between CLST and LILs. The results were visualized using “pheatmap” R package. Comparison of differences between two groups were performed and visualized as box plots or dot plots via “ggplot2” R package and heatmap via “pheatmap” R package according to the guidelines. A *p<0.05 or **p<0.01 was considered statistically significant.

### Diagnostic values evaluation and survival analysis

To test biomarker potential of CLST and LILs for identifying whether CHB patients living with liver injury charactered by abnormal ALT and AST, liver fibrosis, the diagnostic values of immune signals were calculated based on COX analysis using “pROC” R package in liver samples of GSE83148 and GSE84044, respectively. Survival analysis were performed using “survival” R package based on the expression levels of hub genes or ES of the identified immunogenomic signatures in tumor tissues of GSE14520, LIRI-JP and/or CHCC-HBV with available survival data, respectively. Kaplan-Meier curves were draw and plotted via “survminer” R package. LASSO-cox (AI algorithm) for risk model construction and survival prediction was primarily conducted via “survival” R package and “glmnet” R package. A p<0.05 was considered statistically significant.

### AI algorithms for tumor liver samples classification

9 powerful AI algorithms including Logistic Regression (LR), Linear Discriminant Analysis (LDA), K Neighbors (KNN), Gaussian Naive Bayes (GNB), Support Vector Machine (SVM), Random Forest (RF), Decision Tree (CRAT),Gradient Boosting Decision Tree (GBDT) and LightGBM (LGBM, GBDT with leaf-wise) were evaluated for tumor tissues classification. Accuracy (ACC) were calculated as follow:

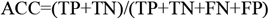

TP: True Positive; FP: False Positive; TN: True Negative; FN: False Negative.

## Results

### Overlapping up-regulated DEGs from mild to severe HBV-LF

Gene expression profiling of HBV-LF were downloaded from GEO database and re-analysis via R studio software according to the previous study(6). As for staging scores, there are 79 up-regulated DEGs in S2 group, 112 up-regulated DEGs in S3 group and 440 up-regulated DEGs in S4 group compared with S0 group, respectively (Fig.1A); Using the online tool Venn 2.1.0, 64 genes that common up-regulated in all of groups from S2 to S4 mentioned above were screened (Fig.1B).

### Functional annotation of overlapping DEGs

As shown in Fig.1C,GO analysis uncovered that DEGs were involved in biological process (BP, 13 items), cellular component (CC, 4 items) and molecular function (MF, 3 items). GO item related to extracellular matrix (CC),a pivotal biological process in HBV-LF was particularly enriched. KEGG signaling pathway analysis revealed that the integrated DEGs were highly enriched in chemokine signaling pathway, cytokine-cytokine interaction, ECM-receptor interaction, Focal adhesion, PI3K-Akt signaling pathway, Toll like receptor pathway and pyrimidine metabolism, respectively (Fig.1D). Among these pathways, chemokine signaling pathway that cargo-carrying genes encoding CXC subfamily ligands (CXCL6, CXCL9, CXCL10, CXCL11) and CCL subfamily ligands (CCL19, CCL20) were primarily enriched.

**Fig 1.**
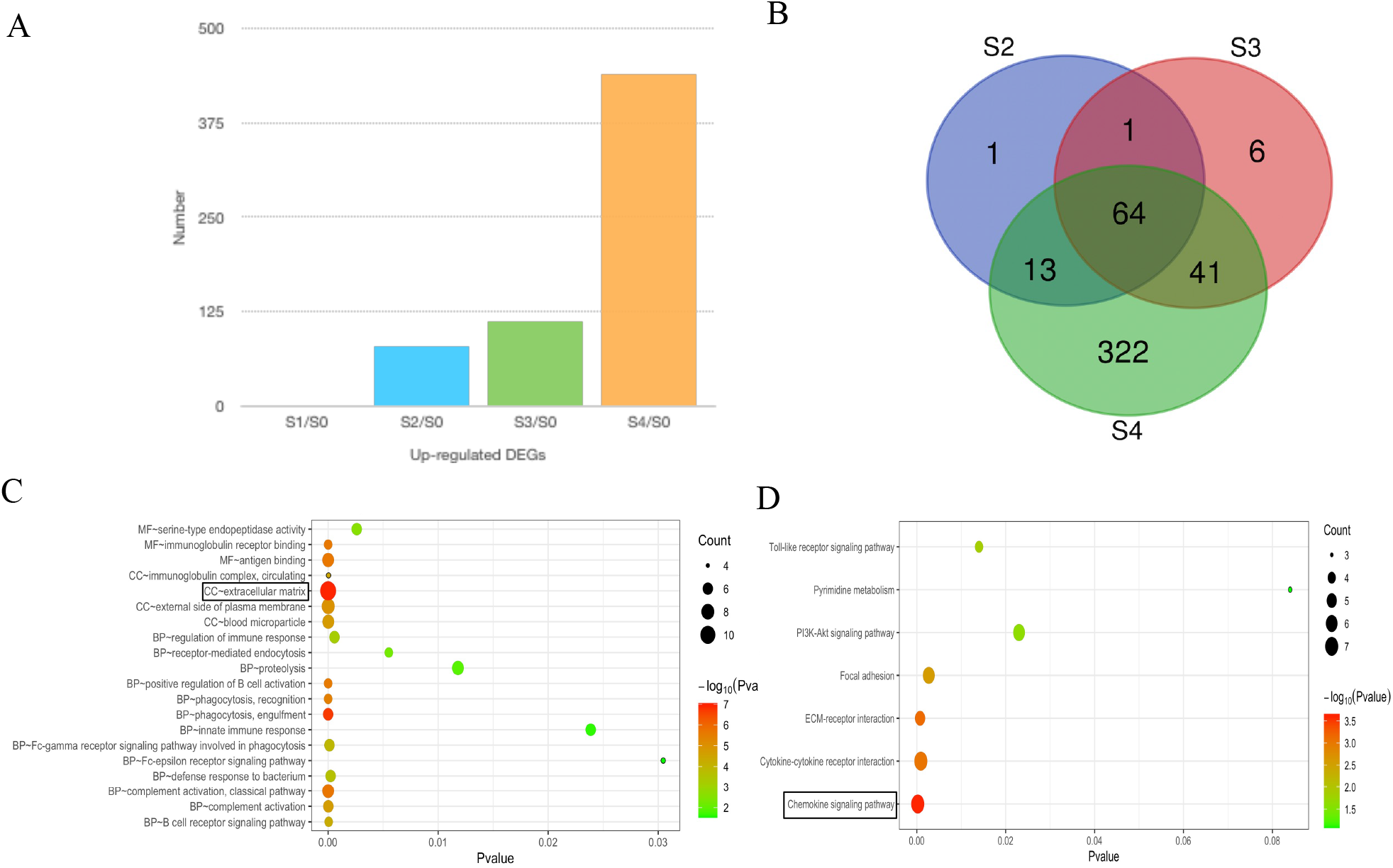
Venn diagram of differentially expressed genes (DEG) and functional annotation in liver fibrosis (GSE84044). (A) Histogram bar chart of up-regulated DEGs at differential staging were drawn by using Numbers. (B) The Venn diagram of the up-regulated DEGs among S2/S0, S3/S0, S4/S0 performed by online tool http://bioinformatics.psb.ugent.be/webtools/Venn/.(C) Visualization of GO analysis of 64 DEGs and extracellular matrix with highest count and lowest p value. MF: molecular function. CC: cellular component; BP: biological process. (D) Visualization of KEGG pathway analysis. Chemokine signaling pathway was highly enriched.

### GS associated hub genes screening

15 hub genes with highest Maximal Clique Centrality (MCC) score including chemokines-related gene cluster (CXCL6, CXCL9, CXCL10, CXCL11, CXCR4, CCL19, CCL20) and ECM-related gene cluster (COL1A1, COL1A2, SPP1, VCAN, LUM, SOX9, THBS1, THBS2) were screened out of 64 DEGs using Cytohubba plugin according to the previous studies (Fig.2A, B)(47, 48). The association of 15 hub genes in patients with differential grading scores (G0-G4 phases, GSE84044) were also investigated and almost all of these genes were significantly up-regulated in G2, G3 and G4 group when compared with G0 group, respectively (Fig.2C-E). As shown in Fig.2F, all hub genes positively associated with fibrosis score and inflammation score were listed as GS associated hub genes.

**Fig 2.**
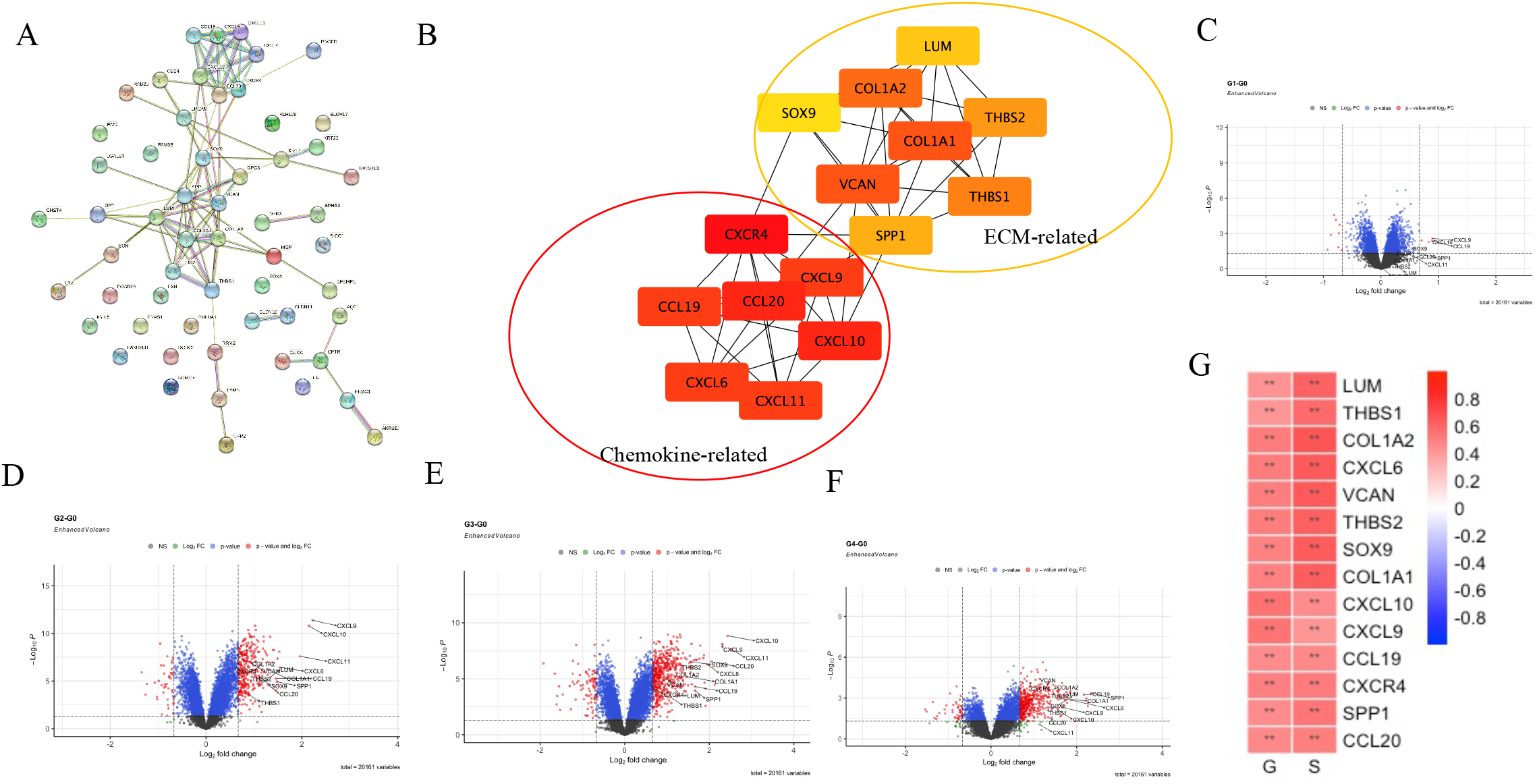
PPI analysis and validation of GS associated hub genes at differential stage of HBV-LF (GSE84044). (A) Original PPI network by STRING. (B) 7 chemokine related genes and 8 ECM related genes in 2 clusters were identified through further evaluation via Cytoscape of MCODE plug-in. (C)-(F) Volcano plots illustrate hub genes that up-regulated in G1, G2, G3 and G4 compared with G0. (G) Correlations between expression levels of hub genes and scores of G or S.

### GS associated hub genes could be induced by initial HBV infection and defined as CLST

GS associated hub genes were explored up-regulated in liver tissues from patients with HBV infection (Fig.3A), abnormal ALT (Fig.3B) and abnormal AST (Fig.3C) in CHB patients. Suppressing, all of these GS associated hub genes were highly enriched in patients at immune clearance phases (IA, immune active) charactered by positive serum HBeAg, unnormal ALT (>40 IU/ml), G/S scores of 2-4 (27) and displayed a similar expression pattern among patients at immune tolerance phases (IT) and immune carrier phases (IC) (Fig.3D).Our research suggested that GS associated hub genes are common overlapping DEGs from CHB to LF. To uncover the original inducers of GS associated hub genes, HBV-mice were analyzed and GS associated hub genes that detectable in liver tissues were obviously up-regulated upon HBV infection *in vivo* (Fig.3E). These genes were further defined as a custom gene set CLST, that was significantly expanded in HBV-PHH *ex vivo* when compared with the controls through performing ssGSEA analysis (Fig.3F). Generally, we concluded that GS associated CLST could be directly induced during initial HBV infection, associated with liver injury and the underlying mechanism is worthy of further exploration.

**Fig 3.**
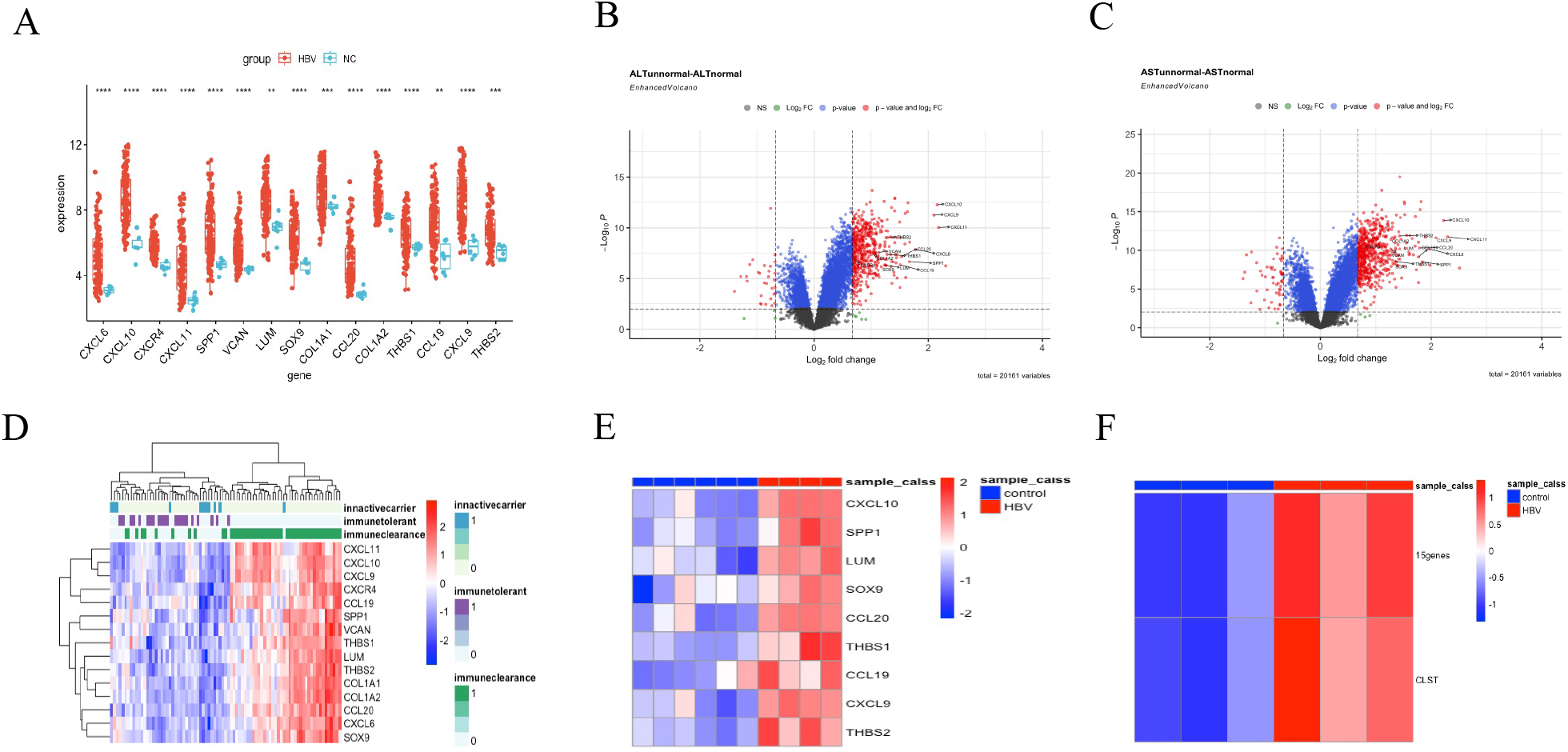
CLST definition. (A) Dot plots of GS associated hub genes in liver samples from CHB group and control group (GSE83148). Blue represents the expression level of hub genes in control group while red represents these genes in liver samples of CHB group. (B)-(C) Volcano plot performed by “EnhancedVolcano” R package showing GS associated hub genes were up-regulated in HBV patients with abnormal ALT and AST when compared with normal ALT and AST, respectively (GSE83148). (D) Heatmap of GS associated hub genes in CHB patients at IC, IT and IA phases (GSE65359). (E) Heatmap of 9 hub genes in HBV infected mice and control group (GSE69590). (F) Heatmap showing ES of 15 hub genes and CLST in HBV-PHH and control PHH (GSE138569).

### CLST, co-expanded with LILs, could effectively predict HBV related liver fibrosis

In this section, CLST was uncovered significantly enriched in CHB patients (Fig.4A) along with majority of LILs and graduate increased with the advancement of G (Fig.4B) and S(Fig.4C). 19/28 LILs were highly enriched in CHB patients (Fig.4A). 25/28 LILs were enriched in patients with liver inflammation (Fig.4B), 24/28 LILs were enriched in patients with LF (Fig.4C). Finally, 17 overlapping LILs were selected (Fig.4D) and the fraction of LILs were highly associated with CLST in liver samples from CHB (Fig.4E) and HBV-LF (Fig.4F). Promising prognosis values of CLST, NKT, MDSC and aCD4 in predicting abnormal ALT level with an AUC above 0.85 (Fig.4G) or abnormal AST (Fig.4H) with an AUC above 0.90 when liver samples from patients with normal level used as controls, respectively. Finally, CLST was ranked as top5 with NKT, MDSC and aCD4 that could effectively segregate LF from normal liver samples with a higher AUC above 0.8 (Fig.4I).

**Fig 4.**
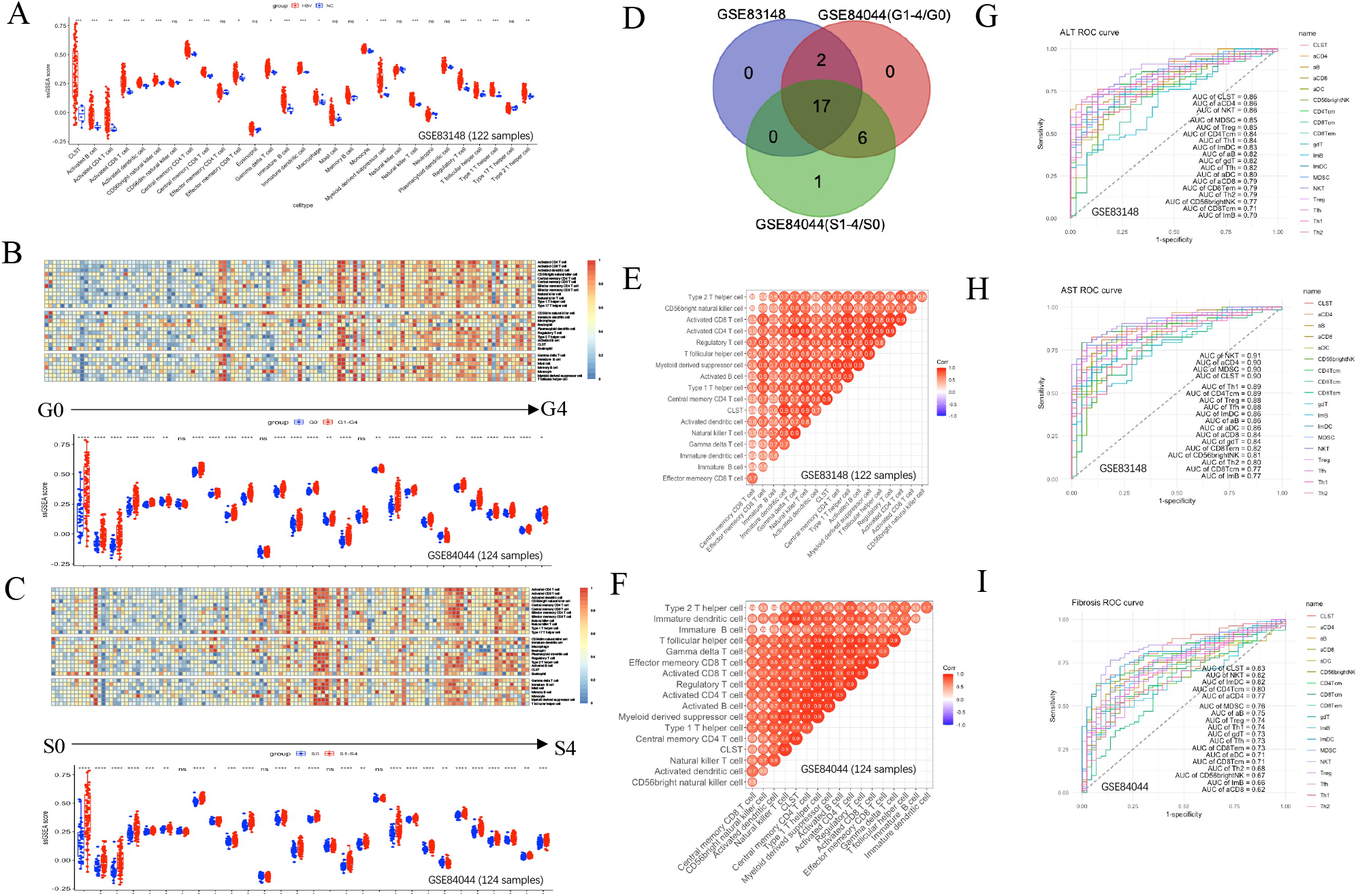
Co-enrichment analysis and diagnostic value of CLST, NKT, MDSC and aCD4 signals. (A) Comparations and boxplot of immune signals between patients with chronic HBV infection and healthy donors (GSE83148). (B) Boxplot of comparisons of immune signals between patients with grading of inflammation(G>=1) and non-inflammation(G=0) (GSE84044). (C) Boxplot of comparisons of immune signals between patients with fibrosis(S>=1) and without fibrosis (S=0). (D) Venn diagram of up-regulated immune signals among GSE83148, GSE84044(S1-4/S0) and GSE84044(G1-4/G0). (E)-(F) Correlation heatmap showing co-enrichment pattern of immune signals in CHB (GSE83148) and in HBV-LF (GSE84044). (G)-(I) ROC curves of immune signals for predicting liver injury and liver fibrosis (GSE84044).

### CLST synergizes with activated CD4+T cells bearing prognostic factors was risk signals in HBV-HCC

There are signs that aCD4+T, NKT and MDSC have certain positive correlations with poor clinical outcome of liver cancer in the original article for 28 LILs definition (31).A correlation analysis were also showed severe positive relationships between CLST and 3 LILs (aCD4, NKT and MDSC) in HBV-HCC cohort with larger sample size of more than 200 patients (Fig.5). TH17 has been widely reported as inflammatory drivers for HCC progression. Recently, liver-resident CD4+T naïve-like cells acquiring a TH17 polarization state has been proved to be a candidate contributor of primary sclerosing cholangitis (PSC) pathogenesis in liver(49). Immune checkpoints(ICs) are involved in poor HCC clinical outcome. To address the question whether CLST and the co-enriched LILs are related to CD4+T_EM_ TH1/TH17 polarization-state and ICs, correlations were examined in tumor tissues from two independent HBV-HCC datasets. Interestingly, obviously positive correlations were observed in tumor tissue from HBV-HCC (Fig.6A). KM analysis showed higher ES of either aCD4+T or CLST was associated with significant shorter OS in HBV-HCC (CHCC, GSE14520) (Fig.6B). Additionally, we investigated member genes of aCD4 by performing PPI analysis and uncovered that CCL20 was the only overlapping gene in CLST and aCD4 (Fig.6C, left). CCL20 was listed as top genes with closed relationship to aCD4 in HBV-HCC (Fig.6C, right). Gene set of aCD4 comprised of 24 genes and 12 genes were explored associated with significant shorter OS in CHCC cohort (Fig6.D). We validated that 6 genes (KIF11, CCNB1, EXO1, KNTC1, PRC1and RGS1) were also risk factors for survival in GSE14520 cohort (data not shown). Either dual higher ES of CLST-aCD4 or aCD4-CCL20 showed a poor survival of HBV-HCC patients (Fig6.E), thus highlights the application of these signals for further prognostic model establishment.

**Fig 5.**
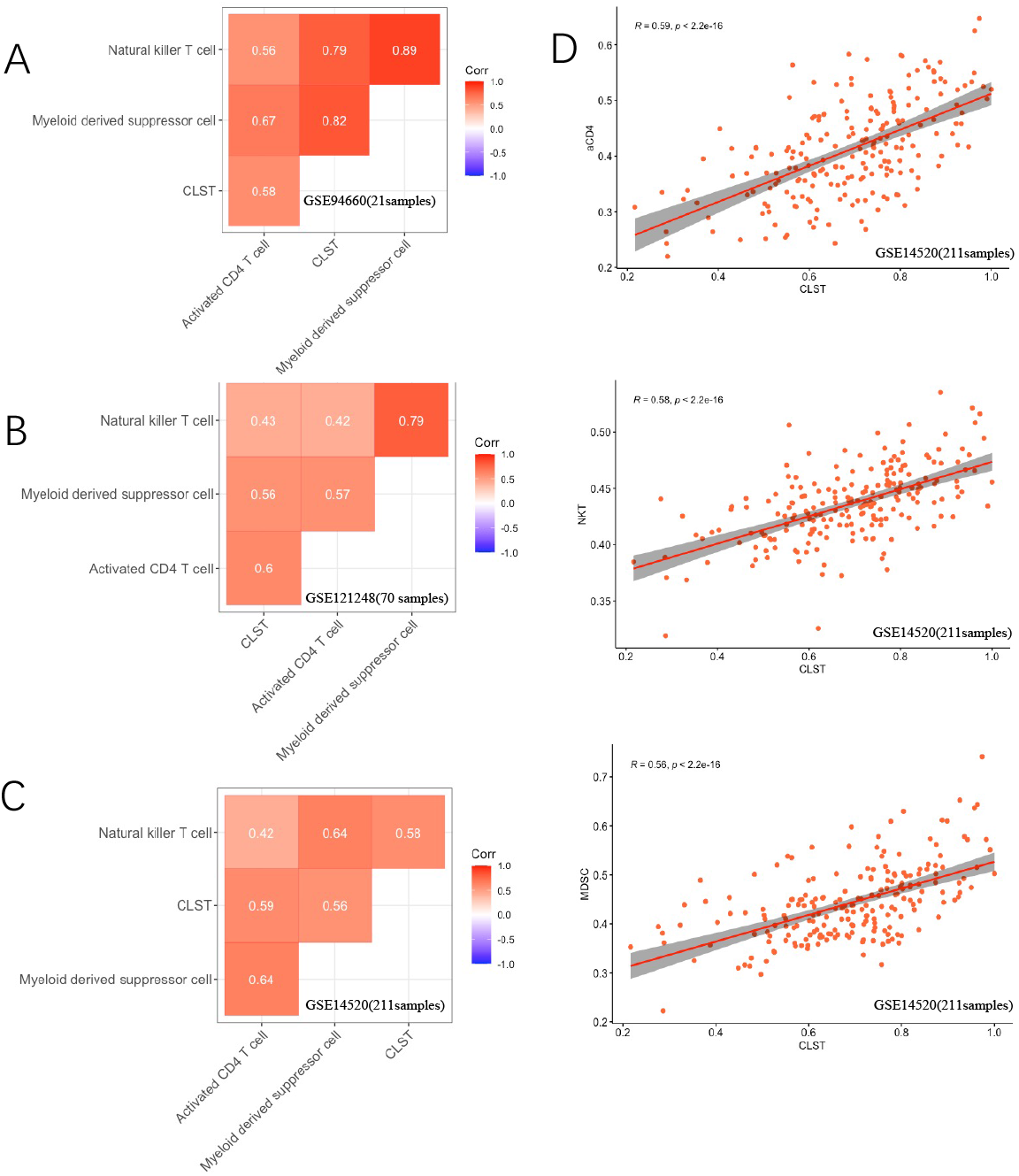
Correlations among CLST and LILs (aCD4, NKT and MDSC) in tumor tissues of HBV-HCC. (A)-(C) Pearson correlation analysis showing co-enrichment among CLST, aCD4,NKT and MDSC signals in three independent GSE datasets (GSE94660, GSE121418 and GSE14520).(D) Pearson correlation matrixes of CLST and LILs (aCD4, NKT and MDSC) in cohort of GSE94660, GSE121418 and GSE14520,respectively.

**Fig 6.**
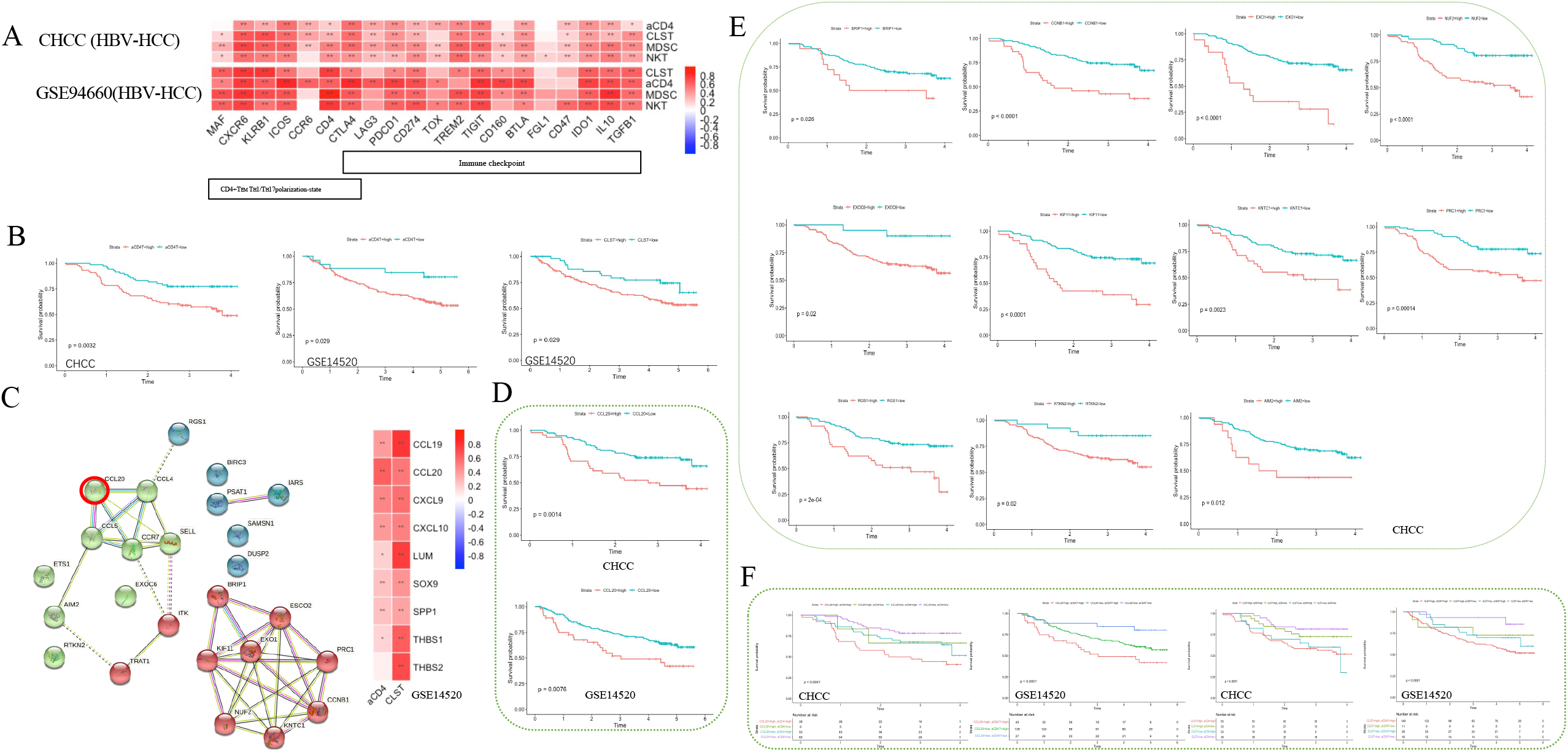
Prognostic factors associated with OS in tumor tissues from HBV-HCC patients. (A) Heatmaps showed correlations between CLST, aCD4, MDSC, NKT signals and selected specific genes in HBV-HCC (CHCC, upper; GSE94660, lower). (B) Kaplan-Meier (KM) Survival plot for patients according to ES of aCD4 and CLST in HBV-HCC (CHCC and GSE14520). (C) PPI analysis of member genes belonging to aCD4 and correlations among ES of aCD4, CLST and expression level of hub genes. CCL20 was an overlapping gene in both aCD4 and CLST. (D) KM analysis of expression level CCL20 in tumor tissues from HBV-HCC patients for OS in two independent cohorts (GSE14520, upper; CHCC, lower). (E) The association between 11 member genes belonging to aCD4 and OS probability in HBV-HCC tumor tissues (CHCC). (F) KM Survival analysis of OS in tumor tissues with dual higher level of aCD4-CCL20 and aCD4-CLST in two datasets (CHCC, GSE14520). Time was calculated by year. Log-rank test for p-value and p-value <0.05 was considered significant.

### Effective prognostic models could be established based on 15 feature genes belonging to CLST and aCD4

15 feature genes comprised of 9 genes (CXCL9, CXCL10, CCL19, CCL20, LUM, SOX9, SPP1, THBS1 and THBS2) from CLST and 6 genes (KIF11, CCNB1, EXO1, KNTC1, PRC1 and RGS1) form aCD4 were ultimate to construct Prognostic Signature Model. Briefly, 9 AI algorithms were trained and validated to separate tumor tissues from healthy liver tissues, cirrhosis tissues and non-tumor tissues (GSE25097). Of 9 algorithms, SVM was performed best in terms of the highest ACC (Fig7.A), showed potent robustness with stratified K fold cross-validations and achieved a high average AUC that can accurately classify tumor tissue from any other types of liver samples (Fig7.B and C). The efficient of SVM was then validated in another independent test set (TCGA-LIHC) with an AUC of 0.97 (Fig7.D). Meanwhile, SVM also showed powerful in predicting patients at early stage (BCLC stage 0-A) of HBV-HCC from non-tumor tissues (GSE14520-HBV-HCC) among all of 9 algorithms (Fig7.E) and achieved an average AUC of 0.99 and 0.99 with stratified K fold cross-validations (splits =5 and 10), respectively (Fig7.F and G). The predictive power of SVM was also excellent in another independent test set (CHCC) with an AUC of 0.99 (Fig8.H). Finally, 15 feature genes and clinical data were ultimate to constructed a prognostic risk model by performing LASSO-COX regression analysis with minimized *λ* (Fig7.I-J). 6 signatures (tumor size, AFP, CCL20, CCNB1, KIF11 and RGS1) risk model was constructed in CHCC cohort. In this model, HBV-HCC patients with high riskScore had a significantly shorter overall survival (Fig 7.K) and AUC score for OS was 0.84 that was higher than single factor (Fig 7.L), respectively.

**Fig 7.**
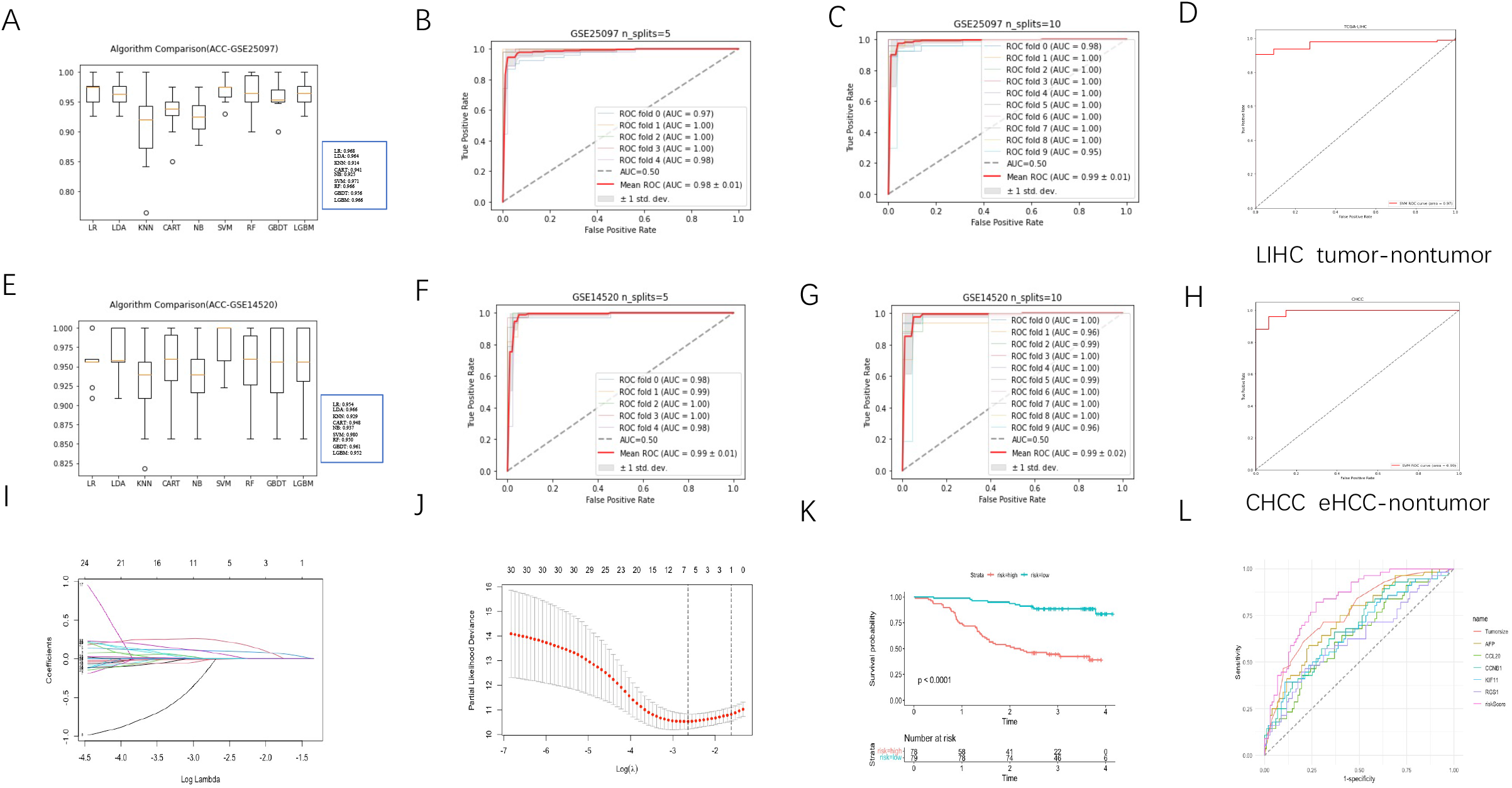
Feature genes belonging to CLST and aCD4 were promising prognostic factors in HBV-HCC. A) Comparison of 11 AI algorithms based on 15 feature genes as a prognostic set with ACC calculation for HCC tumor identification among HCC tumor, adjacent non-tumor, cirrhotic and healthy liver samples. (B)-(C) ROC curves of 15 feature genes for HCC tumor prediction among HCC tumor, adjacent non-tumor, cirrhotic and healthy liver samples via SVM with stratified K fold cross-validations (splices=5 and 10). (D) ROC curves of 15 feature genes as a prognostic set for separation of HCC tumor from non-tumor liver samples via SVM. (E) Comparison of 15 feature genes as a prognostic marker based on 11 AI algorithms with ACC calculation for predicting tumor tissue at early stage of HCC. (F)-(G) The AUC of ROC curves of 15 feature genes as prognostic marker for early stage of HCC tumor separation from non-tumor liver samples via SVM with stratified K Fold cross-validations (splices=5 and 10). (H) ROC curves of 15 feature genes as a prognostic marker for separation tumor tissues at early stage of HCC from non-tumor liver samples via SVM. (I) LASSO coefficient profiles of 30 parameters with k fold cross-validations (k=10). (J) Partial likelihood deviance in LASSO-cox regression model with k fold cross validations(k=10). (K) ROC curves of riskScore, tumor size, AFP, CCL20, CCNB1, KIF11 and RGS1 for predicting OS probability in HBV-HCC (CHCC), respectively.

### IFN-α therapy immune dependently suppressed CLST signal

The landscape of CLST and 28 LILs in liver tissues of pre- and post-IFN-α treatment responders were evaluated. Interestingly, CLST with the infiltration of immune cells tends to be overall down-regulated in these treatment responders (Fig.8A, left). ES of CLST, aCD4 and NKT were significantly suppressed in paired samples (Fig.8A, right) and were also showed significantly positive correlations in these treatment responders (Fig.8B). Distinct expression patterns of 9 hub genes in responders (Fig.8C), HBV-mice *in vivo* (Fig.8D) and HBV-PHHs *ex vivo* (Fig.8E) upon IFN-α treatment was investigated and overall down-regulation trend of 9 hub genes was only appeared in liver samples of patients with HBV infection. Thus, we assumed that modulation of CLST mediated by peg IFN-a therapy was competent immune dependent. Interestingly, there is still limitation in IFN-α therapy that only 2 out of 9 genes (CXCL9 and CXCL10) were significantly suppressed in responders (data not shown). A larger sample size and long-term follow-up are worthy of taking into consideration in the future.

**Fig 8.**
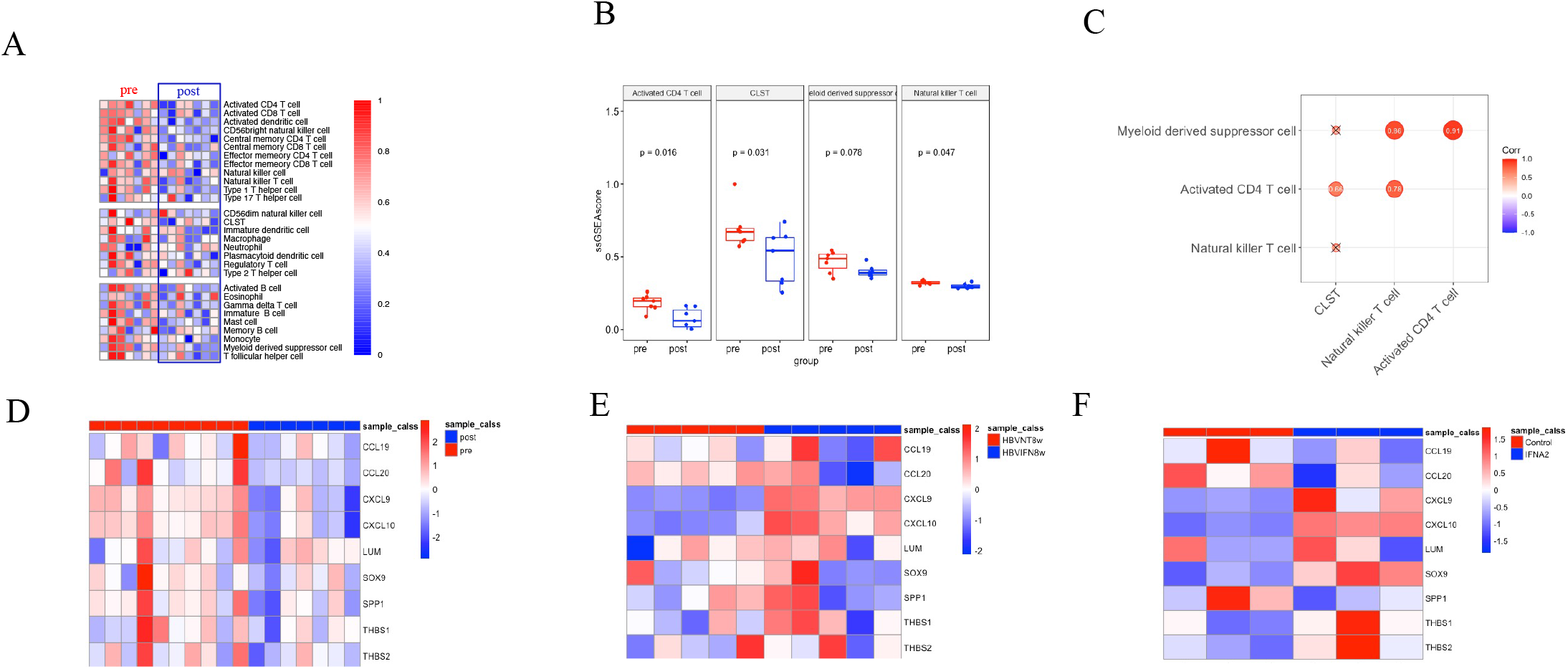
Changes of CLST, LILs and member genes in IFN-α treatment. (A) Heatmaps showed differences between control liver samples and PEG IFN-α treated liver samples (GSE66698). (B) Boxplot of pairwise comparisons of CLST, aCD4, NKT and MDSC between control group and PEG IFN-α treated group (GSE66698). (C) Correlations between CLST, aCD4, NKT and MDSC in PEG IFN-α treated liver samples (GSE66698). (D)-(E) Heatmaps (Upper) showed comparisons of 9 hub genes in control liver samples and treated liver samples (GSE66698, PEG IFN-α treated responders; GSE52752, IFN-α treated HBV-mice; GSE138569, IFN-α treated HBV-PHH).

## Discussion

Chemokine-chemokine receptor and ECM related signals directly and indirectly activated by initial HBV infection are responsible for lymphocytes recruitment, inflammation occurrence, myofibroblast activation, angiogenesis and tumor growth, thus have important impacts on the pathogenesis of HBV related diseases. In this study, 15 hub genes were primarily screened out and observed significantly associated with G and S in a HBV-LF cohort with 124 cases and these genes could be separated into two functional groups(group1 and group2). In detail, members of group 1 are chemokines related genes and of group 2 are ECM related genes. Subsequently,9 of GS associated hub genes were explored up-regulated upon HBV infection in liver tissues of HBV-mice, definite as CLST and confirmed up-regulated in HBV-PHH. We speculate that CLST can be directly induced by HBV infection in hepatocytes at early stage of HBV infection in the absence of immune system *in vivo* and were constantly up-regulated during chronic hepatitis. We also demonstrated that CLST could be served as prognosis signatures for predicting liver injury. These observations suggested that CLST initially induced by HBV should be drivers for HBV pathogenesis and the immune mechanism underlying is worthy of further clarified.

It has been reported that CXCR3-related chemokines (CXCL9 and CXCL10) directly produced by hepatocytes or sinusoidal endothelial cells at early stage of HBV infection can resulted in intrahepatic lymphocyte infiltration (50). The level of intrahepatic mRNA for CXCL10 is remarkably correlated with severity of LF in patients with HIV/HBV co-infection(35) and could promote Th17 recruitment and fuel liver inflammation and LF progression in HBV transgenic (Tg) mice (36). CCL19/CCR7 axis is reported associated with HBV clearance in pAAV/HBV1.2 transfected mice via modulating CD8+T cell responses(51). CCL20 is fibrosis related chemokine in HBV infection (36) and has been shown to be a contributor for HCC initiation and progression through the recruitment of CCR6-positive leukocytes into tumor microenvironment. SPP1(the CD44 ligand) derived by activated HSC serves as a stimulator for KLRG1+ NK cells that can mediate liver scarring limitation in CHB pathogenesis (52) and shows promising prognostic value in HCC (53, 54). SOX9 that can be directly induced in HBV infected human hepatoma cells (55) has been identified as a risk factor in cirrhosis and HCC(56, 57). Serum THBS1 is revealed as a biomarker by proteomic in HBV-LF(58). THBS1 and THBS2 along with pigment epithelium-derived factor (PEDF), have been recently reported involved in angiogenesis and lymph angiogenesis in intrahepatic cholangiocarcinoma(59). Together, majority of member genes of CLST are closely related to LILs and functional transition of LILs in liver microenvironment. However, these previous studies are performed based on flow cytometry(FCM), immune fluorescence (IF) and immunohistochemistry (IHC) with a limited subpopulations of LILs and samples size (50) (36) (51) (52),the cross-talking between CLST with a variety of LILs/immunogenomic pathways are not comprehensively investigated due to the absence of efficient tools for defining further richness of LILs and immunogenomic pathways from high through transcriptomic data until GSEA based algorithms(31, 60) and other algorithms including CIBERSORT(61), MCP-counter, TIMER *et al* were developed(31, 60, 62, 63).Nowadays, GSEA based algorithms have been widely used in tumor(29, 31, 64-67) and non-tumor researches (32, 68, 69) for immunogenomic signatures/pathways identification. According to the previous studies(31, 32, 67, 69),we calculated ES of 28 LILs by cell marker gene based ssGSEA algorithm in HBV related diseases. A comprehensive view of LILs was provided and 21 out of 28 LILS showed closed correlations with CLST in liver tissues from CHB patients and several cell types including MDSC(70-73) and Treg(74-76) previously reported as risk factors during progressive HBV related diseases. Moreover, CLST and the member genes were specifically enriched in patients at IA phases with inadequate immune clearance mediated liver damage when compared with those patients at any other phases. These results partially fulfil the suggestion that host immune responses in liver microenvironment of CHB patients are unfavorable and CLST signal is participant in this disbalanced process.

Although most of hub genes including intrahepatic mRNA for CXCL9 (6, 77), CXCL10 (6, 35, 77), CCL20(36),SOX9(55, 78), SPP1(54), LUM(78) mentioned above have been involved in HBV-LF, there are still rare studies systemically describing the landscape of these genes at whole phages of HBV pathogenesis, even less the integration of these genes as a gene set for HBV-LF predicting. Several studies previously based on bioinformatics has given rise to the evidence that host encoding genes can be served as prognosis biomarkers like CD24, EHF, LUM, SOX9 and ITGBL1 and established Fibrosis Risk Score (FRS) based on these genes for predicting HBV-LF (22, 26, 78), but none of these studies take global immunogenomic information into consideration. In the current study, we uncovered CLST signal was co-enriched with majority of TILs in patients with HBV-LF. CLST, aCD4+T, NKT and MDSC were 4 overlapping metagenes and CLST was ranked as the leading risk factor for separating patients with HBV-LF from those without LF that was comparable to FRS model?

To address the question whether CLST, aCD4, NKT and MDSC are also risk signals in HBV-HCC, 4 HCC cohorts were collected and re-analyzed. Interestingly, co-enrichment among CLST, aCD4, NKT and MDSC were uncovered. Mechanically, CLST, NKT, MDSC and aCD4 were explored highly associated with both CD4+T_EM_ Th1/Th17 polarization-state and levels of ICPs in tumor tissues from patients with HBV-HCC. Th17 recruited via CCL20-CCR6 axis in tumor microenvironment (TME) are contributors for tumorigenesis and drivers of worse clinical outcome (79-82) and ICPs have been well demonstrated account for immunosuppressive microenvironment formation that favor anti-tumor immune evasion (83). These observations give rise to the hypothesis that CLST and LILs may be one of leading causes for unfavorable immune surveillance and TME generation. Correspondingly, either higher ES of CLST or aCD4 implicates shorter OS. Mechanically, we provide insights into the 24 member genes of aCD4 and highlights half of these genes were significantly associated with worse prognosis in HBV-HCC. Interestingly, CCL20 that correlated with worse patient survival rates in HBV-HCC was the only overlapping gene among 33 marker genes specific to CLST and aCD4. Dual higher level of CCL20-aCD4 and CLST-aCD4 were critical for poor progression of HBV-HCC in our further research implying the potential roles of these interaction in immune escape elicitation. Obviously, activated CD4+T cells could be referring to a special CD4+T cell subsets (Th17, Treg., et al) at the station of activation. Nowadays, a verity of novel immune subsets at single-cell level have been detected (84) and the definition of aCD4 in our study bearing CCL20 needs further exploration.

In this study,15 feature genes from CLST and aCD4 were incorporated to perform 9 AI algorithms with cross-validation for detecting tumor tissue at early stage of HBV-HCC from non-tumor tissues, respectively. 15 feature genes based SVM-derived model was built and worked robustly with high accuracy and powerful AUC in both training sets and independent validation sets. We also develop a survival-sensitive risk model holds considerable predative values on patients’ survival through LASSO-COX regression analysis for predicting OS of HBV-HCC patients by comprehensively integrating multiparameter including clinal information, laboratory parameters and feature genes. It is an urgent need for robust tools and workflow to detect early stage of HCC and predict HCC-related death due to the limitation of efficient HCC treatments and these AI models developed in our study is worthy of further development that will be useful at the improvement of clinical decision-making in future anti-tumor therapies. Our bioinformatic analysis of web-accessible data above indicated that CLST was potent signals across all stages of HBV infection from initial HB V infection, CHB, HBV-LF to HBV-HCC. Peg IFN-α treatment in responders has been proved efficient in liver function re-establishment and prevention of HBV-LF and HBV-HCC. Finally, liver transcriptomes of HBV patients before and post standard peg IFN-α treatment were analyzed to test whether first-line therapy exerts anti-HBV effect through modulating CLST signal. Surprisingly, CLST signal was significantly suppressed in responders provide solid evidence of the pivotal roles of this risk signal playing in HBV pathogenesis and the achievement of HBV related diseases cure.

